# An integrative study of five biological clocks in somatic and mental health

**DOI:** 10.1101/2020.06.11.146498

**Authors:** Rick Jansen, Josine Verhoeven, Laura KM Han, Karolina A Aberg, Edwin CGJ van den Oord, Yuri Milaneschi, Brenda WJH Penninx

## Abstract

Biological clocks have been developed at different molecular levels and were found to be more advanced in the presence of somatic illnesses and mental disorders. However, it is unclear whether different biological clocks reflect similar aging processes and determinants. In ~3000 subjects, we examined whether 5 biological clocks (telomere length, epigenetic, transcriptomic, proteomic and metabolomic clocks) were interrelated and associated to somatic and mental health determinants. Correlations between biological clocks were small (all *r*<0.2), indicating little overlap. The most consistent associations with the advanced biological clocks were found for male sex, higher BMI, metabolic syndrome, smoking and depression. As compared to the individual clocks, a composite index of all five clocks showed most pronounced associations with health determinants. The large effect sizes of the composite index and the low correlation between biological clocks, indicate that one’s biological age is best reflected by combining aging measures from multiple cellular levels.

## INTRODUCTION

Aging can be conceptualized in different ways. While chronological age is measured by date of birth, biological age reflects the relative aging of an individual’s physiological condition. Biological aging can be estimated by various cellular indices^1^. Commonly used indices are based on telomere length, DNA methylation patterns (epigenetic age), variation in transcription (transcriptomic age) as well as alterations in the metabolome (metabolomic age) and in the proteome (proteomic age) (see Han et al.^2^, Xia et al.^3^ and Jylhava et al.^4^ for recent reviews). Biological clocks are computed as the residuals of regressing biological age on chronological age: a positive value means that the biological age is larger than the chronological age. Advanced biological aging (i.e an increased biological clock) has been associated to poor somatic health, including the onset of aging-related somatic diseases such as cardiovascular disease, diabetes and cognitive decline^3^. Advanced biological aging has also been correlated to mental health: childhood trauma^5^, psychological stress and psychiatric disorders^6,7^. Specifically, telomere length has been most extensively researched and was found to be shorter in various somatic conditions^8^, all-cause mortality^9,10^ and a range of psychiatric disorders^11^. Advanced epigenetic age has also been linked to worse somatic health, mortality^12^, depressive disorder^7,13^ and post-traumatic stress disorder^14^, although some studies have found associations with the opposite direction of effect^15,16^. Advanced transcriptomic age was found in those with higher blood pressure, cholesterol levels, fasting glucose, and body mass index (BMI)^17^. Advanced metabolomic age increases risk on future cardiovascular disease, mortality, and functionality^18^.

While all biological clocks aim to measure the biological aging process, there is limited evidence for cross-correlations among different clocks. Belsky and colleagues^19^ recently showed low agreement between eleven quantifications of biological aging including telomere length, epigenetic aging and biomarker-composites. In contrast, Hasting and colleagues^20^ showed relatively strong correlations (r>.50) between three physiological composite biological clocks (i.e. homeostatic dysregulation, Klemer and Doubal’s method and Levine’s method), but not with telomere length. Other studies showed that telomere length was not correlated with epigenetic age^7,21^, although cell type composition adjustments revealed a modest association^22^. Further, both Hannum and Horvath epigenetic clocks^23,24^ showed modest correlations to a transcriptomic clock^17^.

Most previous studies, however, have separately considered the relation between single biological clocks and different somatic and mental health conditions. To date, extensive integrated analyses across multiple cellular and molecular aging markers in one study are lacking and it remains unknown to what extent different biological clocks are similarly associated to different health determinants. In addition, most studies did not examine health in its full range and, consequently, whether both somatic and mental health are associated with biological aging remains elusive. As it is unlikely that a single biological clock can fully capture the complexity of the aging process^25^, a composite index, that integrates the different biological clocks and thereby aging at several molecular levels, may reveal the strongest health impact. Therefore, there is an additional need to integrate different biological clocks and test whether such a “composite clock” outperforms single biological blocks in its association with health determinants.

To develop a better understanding of the mechanisms underlying biological aging, this study aimed to examine 1) the intercorrelations between biological clocks based on different molecular levels ranging from DNA to metabolites, namely telomere length, epigenetic, transcriptomic, proteomic and metabolomic clocks; 2) the relationships between different biological clocks with both somatic and mental health determinants; and 3) whether a composite biological clock outperforms single biological clocks in its association with health. For the five biological clocks and the composite clock, associations were computed with a wide panel of lifestyle (e.g. alcohol use, physical activity, smoking), somatic health (functional indicators, BMI, metabolic syndrome, chronic diseases) and mental health (childhood trauma, depression status) determinants.

## RESULTS

### Sample characteristics

To create markers for biological aging we used whole blood derived measurements from the Netherlands Study of Depression and Anxiety (NESDA) baseline assessment: telomere length (*N*=2936), epigenetics (DNA methylation, *N*=1130, MBD-seq, 28M CpGs), gene expression (*N*= 1990, Affymetrix U219 micro arrays, >20K genes), proteomics (*N*=1837, Myriad RBM DiscoveryMAP 250+, 171 proteins) and metabolites (*N*=2910, Nightingale Health platform, 231 metabolites), with 653 overlapping samples (see Table 1 for sample characteristics). Each subsample included around 66% female, with mean age of around 42 years.

**Table 1.**
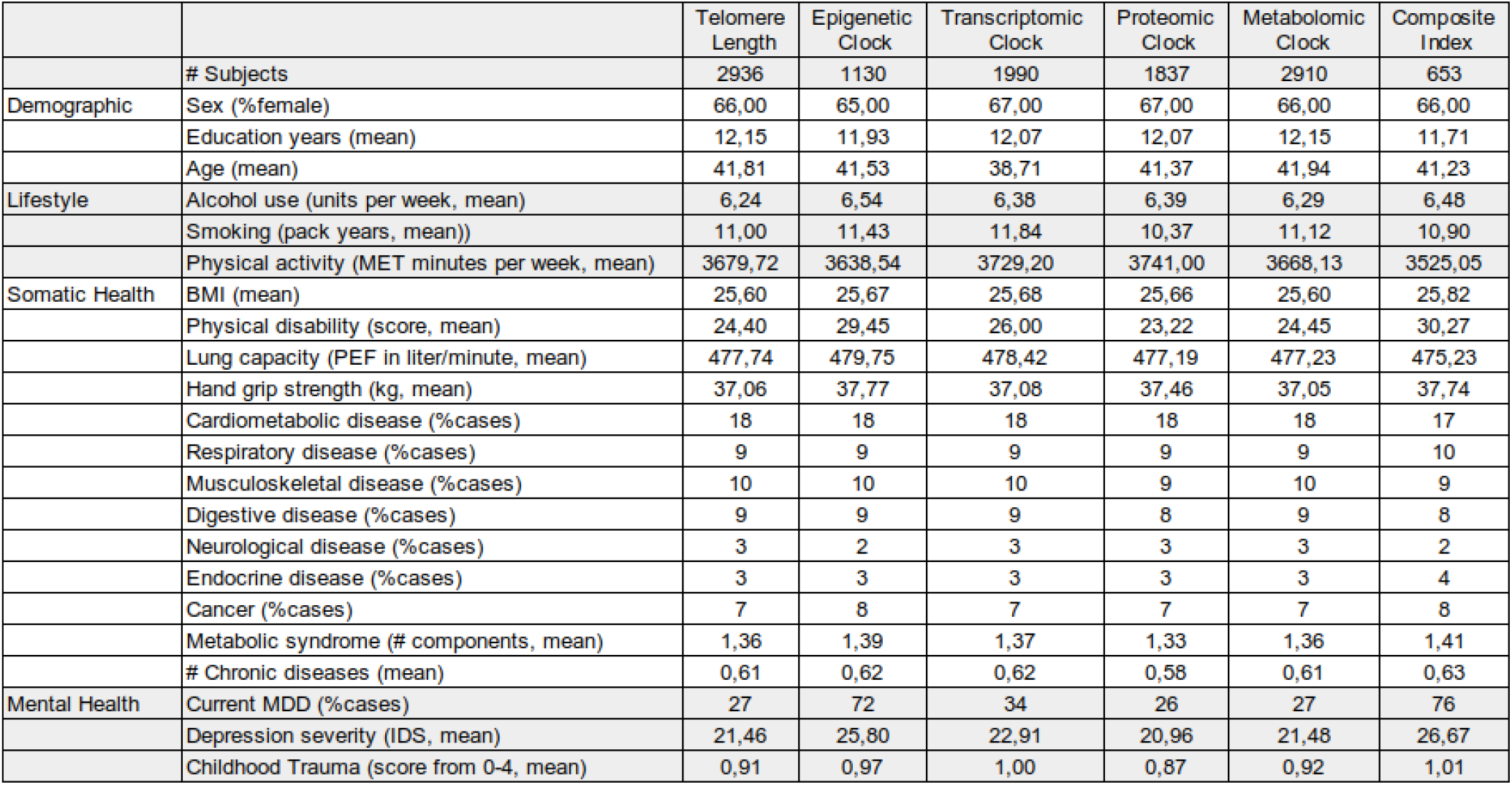
Sample description.

### Computing biological clocks

The methods for creating the biological clocks are describe in detail in the methods section. In brief, for each of the four omic measures (epigenetic, transcriptomic, metabolomic and proteomic) we computed biological age using ridge regression and cross validation (see Figure 1 for study design). Telomere length was multiplied by −1 to be able to compare directions of effects consistent with that of other biological clocks. Correlations between chronological age and biological age markers were 0.30 for telomere length, 0.95 for epigenetic age, 0.72 for transcriptomic age, 0.85 for proteomic age, and 0.70 for metabolomic age (Figure 1). For each omics measure, biological clocks were computed as the residuals of regressing biological age on chronological age. Correlations between biological clocks, corrected for sex, are presented in Figure 2. Correlations were significant for 3 out of 10 pairs; proteomic vs metabolomic clocks (*r*=0.19, *P*=2e-16), transcriptomic vs epigenetic clocks (*r*=0.15, *P*=3e-06) and transcriptomic vs proteomic clocks (*r*=0.08, *P*=2e-06).

**Figure 1.**
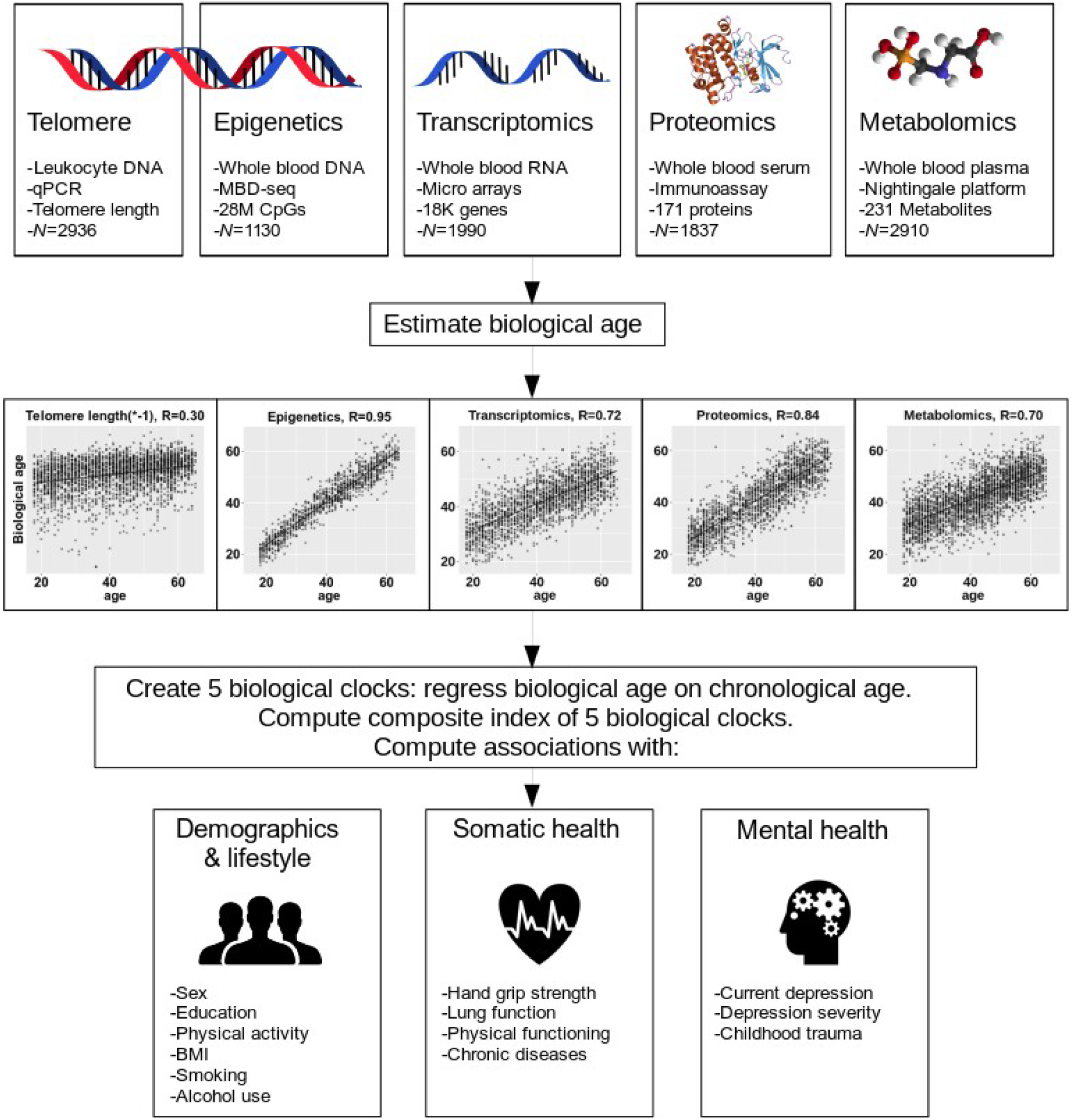
Study Design. The upper part of the figure shows the 5 biological layers. For each of the 5 layers, biological age was estimated, and regressed on age to obtain 5 biological clocks. For the 5 biological clocks associations were computed with multiple demographic, lifestyle, somatic health and mental health determinants.

**Figure 2.**
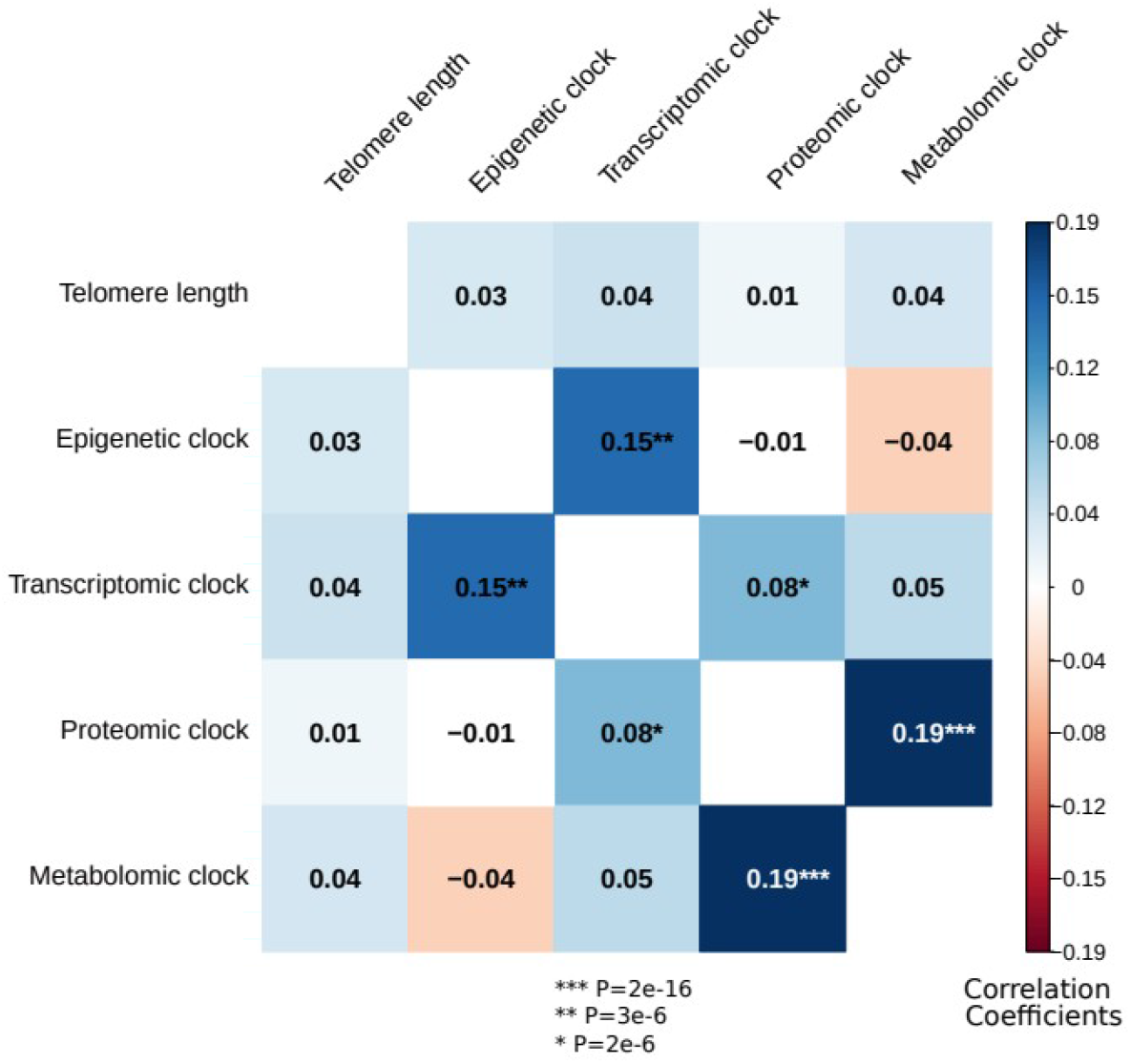
Correlations between the biological clocks. The heatmap represents Spearman rank correlations between the 5 biological clocks, all corrected for sex. Out of ten pairs, three are significant: transcriptomic vs epigenetic clocks, metabolomic vs proteomic clocks and proteomic vs transcriptomic clocks

### Associations between individual biological clocks and health determinants

For each of the five biological clocks we computed associations with several demographic (sex, education), lifestyle (physical activity, smoking, alcohol use), somatic health (BMI, hand grip strength, lung function, physical disability, chronic diseases) and mental health (current depression, depression severity, childhood trauma) determinants. Except for the proteomic clock, sex was associated with all biological clocks: women were biologically younger than men (*P*=3e-4 for telomere length, *P*=5e-4 for the epigenetic clock, *P*=4e-11 for the transcriptomic clock, *P*=1e-5 for the metabolomic clock). Education was not associated with any biological clock. We controlled for sex by using it as a covariate in all following models (only not in the model where sex was the outcome). Table 2 and Figure 3 give an overview of all associations. Correction for multiple testing was done using permutation based FDR (Methods), resulting in a *P*-value threshold of 2e-2 for a FDR of 5% for all tests.

**Table 2.** Associations between 5 biological clocks and multiple health determinants. For each biological clock linear models were fit with the health determinant as predictor, while controlling for sex. Beta’s and P-values from these models are presented here. In the 653 samples with all 5 data layers available, a composite index was constructed which was significantly associated with more variables than any of the 5 biological clocks individually.

|  | Telomere Length | N=2936 | Epigenetic clock | N=1130 | Transcriptomic Clock | N= 1990 | Proteomic Clock | N=1837 | Metabolomic Clock | N=2910 | Composite Index | N=653 |
| --- | --- | --- | --- | --- | --- | --- | --- | --- | --- | --- | --- | --- |
|  | Beta* | P | Beta | P | Beta | P | Beta | P | Beta | P | Beta | P |
| Sex (male/female) | -0.06 | **2,89E-04** | -0.10 | **4,65E-04** | -0.15 | **3,64E-11** | -0.03 | 1,46E-01 | -0.08 | **1,25E-05** | -0.18 | **2,33E-06** |
| Education (# years) | -0.03 | 1,12E-01 | -0.02 | 5,21E-01 | -0.01 | 6,37E-01 | -0.05 | 3,43E-02 | -0.03 | 8,22E-02 | -0.04 | 3,11E-01 |
| Alcohol use (units per week) | 0.03 | 1,05E-01 | -0.05 | 1,40E-01 | 0.00 | 9,21E-01 | 0.07 | **2,89E-03** | 0.04 | 4,57E-02 | 0.07 | 6,05E-02 |
| Smoking (pack years) | 0.06 | **3,11E-03** | 0.02 | 6,22E-01 | 0.05 | **1,55E-02** | 0.10 | **1,33E-05** | 0.05 | **5,09E-03** | 0.10 | **1,15E-02** |
| Physical activity | 0.02 | 2,75E-01 | -0.06 | 3,88E-02 | -0.04 | 6,42E-02 | 0.03 | 1,51E-01 | 0.01 | 5,18E-01 | -0.04 | 3,62E-01 |
| BMI | 0.04 | **1,80E-02** | 0.09 | **3,94E-03** | 0.14 | **6,02E-10** | 0.12 | **9,82E-08** | 0.23 | **2,07E-35** | 0.24 | **2,32E-10** |
| Physical disability | 0.03 | 9,11E-02 | 0.11 | **1,41E-04** | 0.04 | 8,61E-02 | 0.04 | 7,42E-02 | -0.01 | 4,24E-01 | 0.10 | **7,38E-03** |
| Lung capacity | 0.02 | 4,19E-01 | 0.03 | 4,65E-01 | 0.04 | 2,13E-01 | -0.04 | 1,51E-01 | 0.03 | 2,37E-01 | 0.03 | 5,34E-01 |
| Hand grip strength | -0.02 | 3,33E-01 | -0.06 | 1,71E-01 | 0.03 | 3,52E-01 | 0.01 | 7,30E-01 | 0.03 | 2,24E-01 | -0.03 | 6,14E-01 |
| Cardiometabolic disease (no/yes) | 0.02 | 3,37E-01 | 0.04 | 1,56E-01 | 0.03 | 1,44E-01 | 0.03 | 1,35E-01 | 0.05 | **3,94E-03** | 0.10 | **1,37E-02** |
| Respiratory disease (no/yes) | -0.02 | 2,12E-01 | -0.01 | 6,34E-01 | 0.02 | 2,85E-01 | 0.03 | 1,27E-01 | 0.01 | 4,67E-01 | -0.03 | 4,70E-01 |
| Musculoskeletal disease (no/yes) | 0.00 | 8,11E-01 | -0.01 | 7,37E-01 | 0.04 | 1,04E-01 | 0.02 | 4,36E-01 | 0.02 | 2,23E-01 | 0.09 | 2,27E-02 |
| Digestive disease (no/yes) | 0.03 | 5,77E-02 | -0.02 | 5,71E-01 | 0.06 | **9,76E-03** | 0.06 | **1,21E-02** | 0.02 | 2,81E-01 | 0.05 | 2,01E-01 |
| Neurological disease (no/yes) | -0.02 | 2,58E-01 | 0.02 | 5,60E-01 | 0.01 | 5,44E-01 | 0.02 | 2,84E-01 | 0.02 | 1,93E-01 | -0.04 | 2,64E-01 |
| Endocrine disease (no/yes) | -0.01 | 4,45E-01 | 0.01 | 8,13E-01 | -0.01 | 5,75E-01 | 0.06 | **1,03E-02** | 0.03 | 1,23E-01 | 0.06 | 1,18E-01 |
| Cancer (no/yes) | 0.00 | 9,66E-01 | 0.02 | 5,65E-01 | 0.02 | 4,88E-01 | 0.03 | 1,81E-01 | 0.02 | 2,01E-01 | 0.08 | 3,22E-02 |
| Metabolic syndrome (# components) | 0.06 | **6,35E-04** | 0.04 | 1,46E-01 | 0.13 | 9,98E-09 | 0.13 | **5,34E-09** | 0.21 | **4,53E-29** | 0.28 | **9,10E-13** |
| # Chronic diseases | 0.00 | 7,99E-01 | 0.03 | 3,63E-01 | 0.05 | 3,20E-02 | 0.09 | **1,24E-04** | 0.03 | 1,39E-01 | 0.06 | 1,26E-01 |
| Current MDD (no/yes) | 0.03 | 1,59E-01 | 0.09 | **1,99E-03** | 0.07 | **1,68E-02** | 0.08 | **7,62E-03** | -0.03 | 1,61E-01 | 0.11 | **6,05E-03** |
| Depression severity | 0.04 | 2,40E-02 | 0.12 | **8,67E-05** | 0.03 | 2,76E-01 | 0.07 | **5,99E-03** | -0.02 | 3,74E-01 | 0.13 | **7,61E-04** |
| Childhood Trauma | 0.01 | 4,54E-01 | 0.12 | **7,99E-05** | 0.03 | 2,06E-01 | 0.04 | 8,96E-02 | 0.04 | 2,46E-02 | 0.09 | **1,96E-02** |

**Figure 3.**
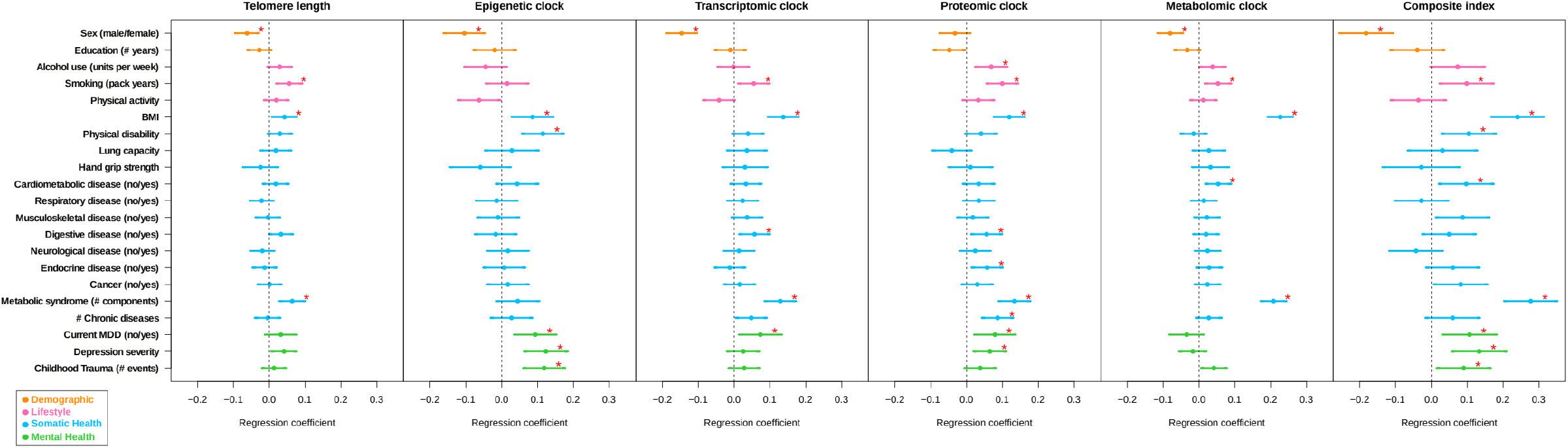
Forest plot of associations between biological clocks and health determinants. For each of the associations between biological clocks and health determinants, the standardized beta and standard deviation derived from linear models were plotted. The significant associations (P<2e-2, FDR<5%) are shown with red stars. The composite index, which is the scaled sum of the 5 biological clocks, clearly shows most associations and often largest effect sizes.

Among the lifestyle indicators, alcohol use was associated with an advanced proteomic clock (*P*=3e-3) and smoking (packs per year) was associated with shorter telomere length (*P*=3e-3), and advanced transcriptomic (*P*=2e-2), proteomic (*P*=1e-5) and metabolomic clocks (*P*=5e-3). Physical activity was not associated with any biological clock.

From the somatic health determinants, high BMI was strongly associated with an advancement of all biological clocks (*P*=2e-2 for telomere length, *P*=4e-3 for the epigenetic clock, *P*=6e-10 for the transcriptomic clock, *P*=1e-7 for the proteomic clock, and *P*=2e-35 for the metabolomic clock). Physical disability was associated with an advanced epigenetic clock (*P*=1e-4). Within the domain of chronic diseases, the presence of digestive disease and endocrine disease were associated with an advanced proteomic clock (*P*=2e-2 and *P*=1e-2, respectively). Subjects with cardiometabolic disease had an advanced metabolomic clock (*P*=4e-3) and subjects with digestive disease had an advanced transcriptomic clock (*P*=1e-2). Those with metabolic syndrome showed four advanced biological clocks (*P*=6e-4 for telomere length, *P*=1e-8 for the transcriptomic clock, *P*=5e-9 for the proteomic clock, *P*=5e-29 for the metabolomic clock).

The presence of current depression and depression severity were associated with advanced epigenetic (*P*=2e-3 and *P*=9e-5) and proteomic (*P*=8e-3 and *P=6*e-3 respectively) clocks. Current depression was also associated with an advanced transcriptomic clock (*P*=2e-2) and those with childhood trauma had an advanced epigenetic clock (*P*=8e-5).

### Associations between the composite index of biological clocks and all health determinants

The composite index was computed as the sum of the 5 scaled biological clocks in the 653 samples with data of all 5 biological levels. Correlations between the 5 biological clocks and the composite index were between 0.43 and 0.51. We found more and stronger associations for the composite index than for any of the individual biological clocks: including sex (*P*=2e-6), BMI (*P*=2e-10), smoking (*P*=2e-2), metabolic syndrome (*P*=9e-13), current MDD (*P*=6e-3), depression severity (*P*=7e-4) and childhood trauma (*P*=2e-2). To allow for direct effect size comparisons, we compared the findings for the composite index to those of each individual biological clock with the same subsample. In this analysis, *P*-values and effect sizes were often more pronounced for the composite index (Figure 4, Table S1). For example, for sex, BMI, metabolic syndrome and current MDD, that were significantly associated with the composite index, the betas for the composite index were larger than the betas from all individual clocks. For the other 5 variables significantly associated with the composite index (smoking, physical disability, cardiometabolic disease, depression severity and childhood trauma) the betas for the composite index were larger than 4 out of 5 betas from the individual clocks.

**Figure 4.**
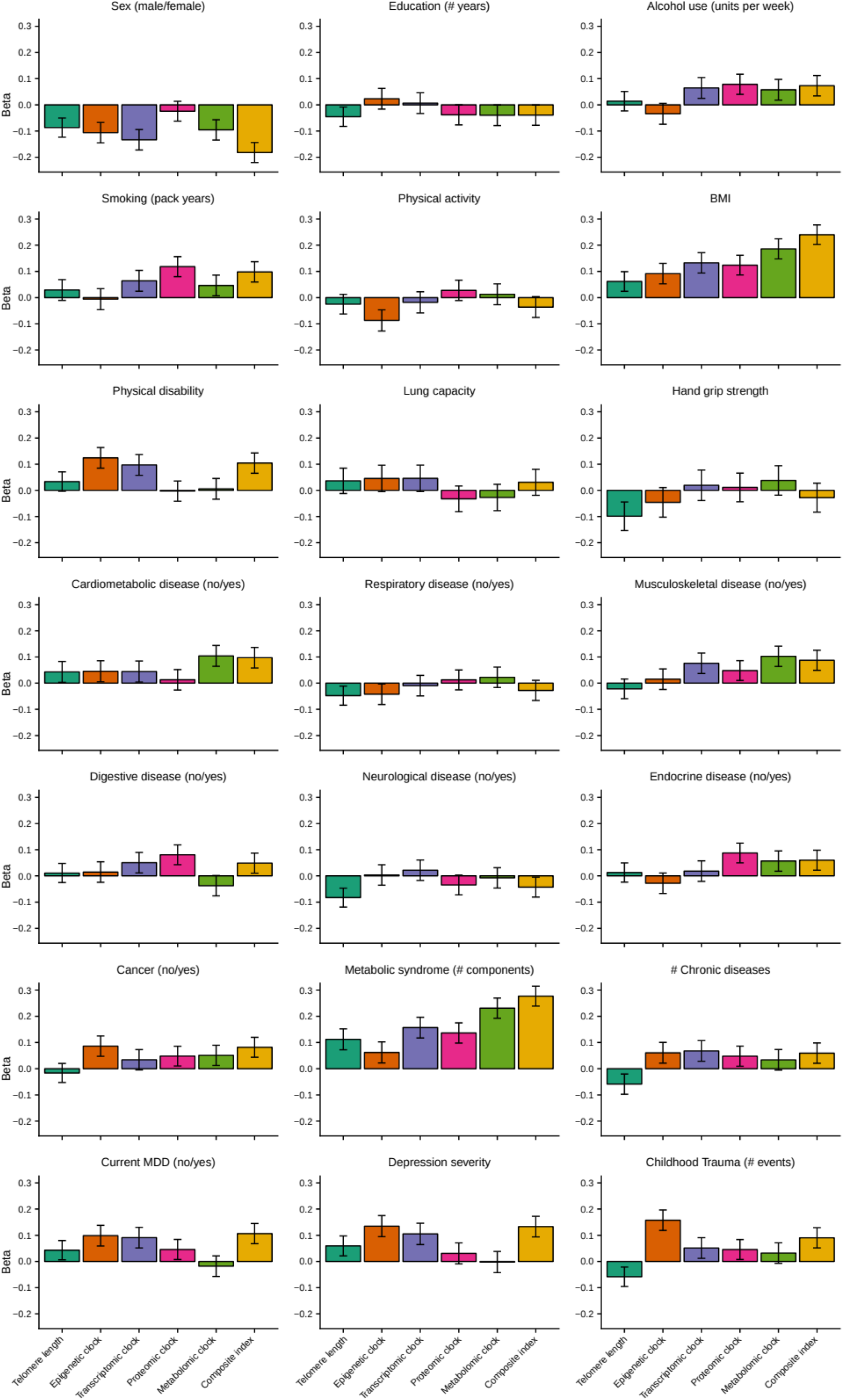
Barplots of betas from associations between biological clocks and health determinants. For each of the associations between biological clocks and health determinants, the standardized beta and standard deviation derived from linear models were plotted. Only samples were used that had data for all 5 biological clocks (N=653).

## DISCUSSION

In this study, we examined five biological clocks based on telomere length and four omics levels from a large, clinically well-characterized cohort. We demonstrated significant intercorrelations between three pairs of biological clocks, illustrating the complex and multifactorial processes of biological aging. Furthermore, we observed both overlapping and unique associations between the individual clocks and different lifestyle, somatic and mental health determinants. Separate linear regressions showed that male sex, high BMI, smoking, and metabolic syndrome were consistently associated with more advanced levels of biological aging across at least four of the biological clocks. Strikingly, depression was associated to more advanced levels of epigenetic, transcriptomic and proteomic clocks, signifying that both somatic and mental health is associated with the biological clocks. Finally, by integrating a composite index of all biological clocks we were able to obtain larger effect sizes with e.g. physical disability and childhood trauma exposure, underscoring the broad impact of determinants on cumulative multi-system biological aging.

Biological aging seems to be differently manifested at certain cellular levels, as suggested by the range of correlations among the biological clocks considered in this study. Consistent with prior studies we showed weak correlations between different biological clocks^26^ and we confirm the absent relationship between telomere length and the epigenetic clock^21,27,28^, but also show lack of associations with the transcriptomic, proteomic or metabolomic clocks. However, we do confirm an earlier finding showing a significant but modest correlation between epigenetic and transcriptomic age^29^. The correlation between the metabolomic and proteomic clocks may partly be explained by the fact that both data were obtained from platforms that were aimed at probing central inflammation lipid processes, rather than the full proteome or metabolome. Nevertheless, we can infer that only some biological clocks show overlap, while most of them seem to be tracking distinctive parts of the aging process, even if they are associated with the same somatic or health determinants.

Our study showed that several of the determinants considered exhibited consistent associations with different biological clocks. First, male sex was associated with shorter telomere length and advanced epigenetic, transcriptomic and metabolomic clocks, in line with a large body of literature that shows advanced biological aging and earlier mortality in males compared to females^30^. Second, high BMI was consistently related to all biological clocks, showing that the more overweight or obese, the higher the biological age^31^, also after controlling for sex. Earlier studies showed similar associations between high BMI and short telomere length^31^, and older epigenetic^32^ and transcriptomic aging signatures^17^. Third, our analyses showed similarly consistent associations between the prevalence of metabolic syndrome and advanced levels of aging. Further, all but the epigenetic clock were advanced with respect to cigarette smoking.

Major depressive disorder (MDD) status was consistently related to advanced aging in three (epigenetic, transcriptomic, proteomic) out of the five biological clocks. In contrast, a recent study (N>1000) in young adults (20-39 years) did not show associations between mental health (as measured by the CIDI) and biological aging (indicated by telomere length, homeostatic dysregulation, Klemer and Doubal’s method and Levine’s method)^20^, but it seems possible that this sample was too young to fully develop aging-related manifestations of mental health problems, or lacked age variation. It is likely that our data (obtained from participants 18-64 years) may have been more sensitive in picking up associations with mental health due to increased variation in both chronological age (i.e. inclusion of older persons), as well as symptom severity.

Furthermore, we computed a composite index by summing up the five biological clocks studied here. In other words, this integrative metric contains cumulative independent signal from the individual markers and dependent shared signal - with possible reduced noise due to the summation - between them. Given that this composite index demonstrates larger effect sizes for BMI, sex, smoking, depression severity, and metabolic syndrome than the individual clocks, it is suggested that being biologically old at multiple cellular levels has a cumulative multi-systemic effect. When integrated, the composite index reveals stronger (i.e. greater cumulative betas for the composite index than individual clocks) converging associations with sex, BMI, metabolic syndrome and current MDD. This provides further support for the hypothesis that not one biological clock sufficiently captures the biological aging process and that not all clocks are under the control of one unitary aging process. There is abundant room for further progress in determining whether biological aging can be modified by intervening on these determinants.

Nonetheless, the question remains which biological mechanism could plausibly link the current quantification of biological aging and its lifestyle, somatic, and mental health determinants. Part of this answer requires discussion on the features used to build the different clocks: the proteomic and metabolomic clocks mostly measure inflammatory or metabolic factors, two highly integrated processes in aging and aging-related diseases^33^. Previous studies suggest immune-mediated mechanisms (specifically inflammatory signaling) connecting metabolic syndrome^34^, mental health disorders^35^, and aging^36^. Moreover, MDD is a condition in which inflammation, obesity, and premature or advanced aging co-occur and converge. It might therefore be speculated that immunity and “inflammaging”^37^ may tie together the currently observed associations.

More research is needed to elucidate whether: 1) physiological disturbances, such as loss of inflammatory control associated with somatic and psychopathology, accelerate biological aging over time, 2) advanced biological aging precedes and constitutes a vulnerability factor that causes somatic and psychopathology, or 3) somatic and psychopathology and biological aging processes are not causally linked, but share underlying etiological roots (e.g. shared genetic risks or environmental factors)^2^. Yet, it could conceivably be hypothesized that dysregulation of immunoinflammatory control may be related to metabolic outcomes, aging, and depression^38^, providing scope as to why some of these determinants converge across different platforms and multiple biological levels.

### Strengths and limitations

Here, we used a large cohort that was well-characterized in terms of demographics, lifestyle, and both somatic and mental health assessments, to study and integrate five biological clocks across multiple levels of analysis. This is particularly important as we show that the determinants of biological aging encompass several different domains. Moreover, our sample was adequately powered to detect statistically significant associations, limiting the possibility for chance findings and increasing probability for identifying robust biological age determinants. On the other hand, an obvious limitation is the cross-sectional nature of this study that prevents us from drawing any conclusions on whether the determinants accelerate the aging trajectory over time, or the other way around. Another limitation is that it is difficult to generalize our prediction models to data from other platforms generating the same datatypes from different probes, emphasizing the need for epidemiological replication of these determinants in other datasets. We recognize that data harmonization and pooling are important strategies on the scientific research agenda that may overcome this limitation in the future.

### Conclusions

In conclusion, this study examined the overlap between five biological clocks and their shared and unique associations with somatic and mental health. Our findings indicate that they largely track distinct, but also partially overlapping aspects of this aging process. Further, we demonstrated that male sex, smoking, higher BMI and metabolic syndrome were consistently related to advanced aging at multiple biological levels. Remarkably, our study also converges evidence of depression and childhood trauma associations across multiple platforms, cellular levels, and sample sizes, highlighting the important link between mental health and biological aging. Taken together, our findings contribute to the understanding and identification of biological age determinants, important to the development of end points for clinical and epidemiological research.

## METHODS

### Study design and participants

Data used were from the Netherlands Study of Depression and Anxiety (NESDA), an ongoing longitudinal cohort study examining course and consequences of depressive and anxiety disorders. The NESDA sample consists of 2981 persons between 18 and 65 years including persons with a current or remitted diagnosis of a depressive and/or anxiety disorder (74%) and healthy controls (26%). Persons with insufficient command of the Dutch language or a primary clinical diagnosis of other severe mental disorders, such as severe substance use disorder or a psychotic disorder were excluded. Participants were recruited between September 2004 and February 2007 and assessed during a 4-hour clinic visit. The study was approved by the Ethical Review Boards of participating centers, and all participants signed informed consent. The population and methods of the NESDA study have been described in more detail elsewhere^39^.

Data to derive different biological clocks was available for different subsamples and all based on a fasting blood draw from participants in the morning between 8:30 and 9:30 after which samples were stored in a −80°C freezer or – for RNA - transferred into PAXgene tubes (Qiagen, Valencia, California, USA) and stored at −20°C. To create biological clocks, we used telomere length (*N*=2936), DNA methylation (*N*=1130, MBD-seq, 28M CpGs), gene expression (*N*=1990, Affymetrix U219 micro arrays, >20K genes), proteomics (*N*=1837, Myriad RBM DiscoveryMAP 250+, 171 proteins) and metabolites (*N*=2910, Brainshake platform, 231 metabolites) which had been measured for other projects already. See Table 1 and details in the following.

### Biological clock assessments

#### Telomere length

Leukocyte telomere length was determined at the laboratory of Telomere Diagnostics, Inc. (Menlo Park, CA, USA), using quantitative polymerase chain reaction (qPCR), adapted from the published original method by Cawthon et al.^40^. Telomere sequence copy number in each patient’s sample (T) was compared to a single-copy gene copy number (S), relative to a reference sample. The resulting T/S ratio is proportional to mean leukocyte telomere length. The detailed method is described elsewhere^41^. The reliability of the assay was adequate: eight included quality control DNA samples on each PCR run illustrated a small intra-assay coefficient of variation (CV=5.1%), and inter-assay CV was also sufficiently low (CV=4.6%).

#### DNA methylation (Epigenetic clock)

To assay the methylation levels of the approximately 28 million common CpG sites in the human genome, we used an optimized protocol for MBD-seq^7,42^. With this method, genomic DNA is first fragmented and the methylated fragments are then bound to the MBD2 protein that has high affinity for methylated DNA. The non-methylated fraction is washed away and only the methylation-enriched fraction is sequenced. This optimized protocol assesses about 94% of the CpGs in the methylome. The sequenced reads were aligned to the reference genome (build hg19/GRCh37) with Bowtie2(32) using local and gapped alignment. Aligned reads were further processed using the RaMWAS Bioconductor package33) to perform quality control and calculate methylation scores for each CpG.

#### Gene expression (Transcriptomic clock)

RNA processing and assaying-done at Rutgers University Cell and DNA repository-have been described previously^27–29^. Samples were hybridized to Affymetrix U219 arrays (Affymetrix, Santa Clara, CA). Array hybridization, washing, staining, and scanning were carried out in an Affymetrix GeneTitan System per the manufacturer’s protocol. Gene expression data were required to pass standard Affymetrix QC metrics (Affymetrix expression console) before further analysis. We excluded from further analysis probes that did not map uniquely to the hg19 (Genome Reference Consortium Human Build 37) reference genome sequence, as well as probes targeting a messenger RNA (mRNA) molecule resulting from transcription of a DNA sequence containing a single nucleotide polymorphism (based on the dbSNP137 common database). After this filtering step, data for analysis remained for 423,201 probes, which was summarized into 44,241 probe sets targeting 18,238 genes. Normalized probe set expression values were obtained using Robust Multi-array Average (RMA) normalization as implemented in the Affymetrix Power Tools software (APT, version 1.12.0, Affymetrix). Data for samples that displayed a low average Pearson correlation with the probe set expression values of other samples, and samples with incorrect sex-chromosome expression were removed.

#### Proteins (Proteomic clock)

As described previously^46^, a panel of 243 analytes (Myriad RBM DiscoveryMAP 250+) involved in various hormonal, immunological, and metabolic pathways was assessed in serum using multiplexed immunoassays in a Clinical Laboratory Improvement Amendments (CLIA)-certified laboratory (Myriad RBM; Austin, TX, USA;). After excluding analytes with more than 30% missing data (mostly due to values outside the ranges of detection), 171 of the 243 analytes remained for analysis (with values below and above detection limits imputed with the detection limit values).

#### Metabolites (Metabolomic clock)

Metabolite measurements have been described in detail previously^18,47^. In short, a total of 232 metabolites or metabolite ratios were reliably quantified from Ethylenediaminetetraacetic acid plasma samples using targeted high-throughput proton Nuclear Magnetic Resonance (^1^ H-NMR) metabolomics (Nightingale Health Ltd, Helsinki, Finland)^48^. Metabolites measures provided by the platform include 1) lipids, fatty acids and low-molecular-weight metabolites (N=51); 2) lipid composition and particle concentration measures of lipoprotein subclasses (N=98); 3) metabolite ratios (N=81). This metabolomics platform has been extensively used in large-scaled epidemiological studies in the field of diabetes, cardiovascular disease, mortality and alcohol intake^18,49–52^. The data contained missing values due to detection limits. Samples with more than 25 missings were removed (N=71), metabolites with more than 250 missings were removed (N=1). Other missing values were replaced with the median value per metabolite. In total 231 metabolites in 2910 samples remained for analysis.

### Building biological clocks for multiple omics domains

Telomere length was multiplied by −1 to be able to compare directions of effects consistent with that of other biological clocks. For each of the other four *omic* domains (epigenetic, transcriptomic, metabolomic and proteomic data) the same approach was used to compute biological clocks. First, the omic data were residualized with respect to technical covariates (batch, lab). Second, data per omics marker were normalized using a quantile-normal transformation. Finally, biological age were computed using cross-validation by splitting the sample in 10 equal parts. For each of the ten groups, 9 parts were used as training set and the 10^th^ as test set. In the training set the biological age estimator was computed using ridge regression (R library glmnet), with chronological age as the outcome, and the omics data as predictor. Only for methylation and gene expression a selection of predictors (CpGs for methylation based models and genes for gene expression based models) was made: CpGs/genes were ranked based on their association with age, using a step wise procedure CpGs/genes were added to the model in the order of their ranks, until the prediction of age did not improve anymore^7^. This resulted in 80,000 CpGs (mapping to 2,976 genes) for the methylation based model, and 1,200 probes (mapping to 767 genes) for the gene expression based model. For the proteomic and metabolomic data, all markers were used to predict age, since leaving markers out decreased the prediction accuracy. The predictor was then used in the test set to create an unbiased omics-based biological age. For each omics domain, biological clocks were computed as the residuals of regressing biological age on chronological age^7,17^. Thus, in the terminology we use here, a biological clock represents the biological age acceleration: a positive value means that the biological age is larger than the chronological age. In the following we will call the biological clocks ‘epigenetic clock’, transcriptomic clock’, etc. A composite index of biological clocks was made by scaling each of the 5 biological clocks and taking the sum.

### Health determinants

#### Lifestyle

Alcohol consumption was assessed as units per week by using the AUDIT^53^. Smoking status was assessed by pack years (smoking duration * cigarettes per day/20). Physical activity^54^ was assessed using the International Physical Activity Questionnaire (IPAQ)^55^ and expressed as overall energy expenditure in Metabolic Equivalent Total (MET) minutes per week (MET level * minutes of activity * events per week).

#### Somatic health

Body mass index (BMI) was calculated as measured weight divided by height-squared. Functional status is one of the most potent health status indicators in predicting adverse outcomes in aging populations^56^, including depression^57^. Assessment of functional status includes measures of physical impairments and disability, reflecting how individuals’ limitations interact with the demands of the environment. Two measures of physical impairments were available: Lung capacity was determined by measuring the peak expiratory flow (PEF in liter/minute) using a mini Wright peak flow meter. Hand grip strength was measured with a Jamar hand held dynamometer in kilograms of force and was assessed for the dominant hand. Furthermore, physical disability was measured with the World Health Organization Disability Assessment Schedule II (WHODAS-II)s the sum of scale 2 (mobility) and scale 3 (self-care). The number of self-reported current somatic diseases for which participants received medical treatment was counted. We used *somatic disease categories* as categorized previously^54,58^: cardiometabolic, respiratory, musculoskeletal, digestive, neurological and endocrine diseases, and cancer. Metabolic syndrome components included waist circumference, systolic blood pressure, HDL cholesterol, triglycerides and glucose levels, which measurement methods are described elsewhere^59^.

#### Mental health

Presence of current (6-month recency) major depressive disorder was assessed by the DSM-IV Composite International Diagnostic Interview (CIDI) version 2.1. Depressive severity levels in the week prior to assessment were measured with the 28-item Inventory of Depressive Symptomatology (IDS) self-report^60^. Childhood trauma was assessed with the Childhood Trauma Interview (CTI)^61^. In this interview, participants were asked whether they were emotionally neglected, psychologically abused, physically abused or sexually abused before the age of 16. The CTI reports the sum of the categories that were scored from 0 to 2 (0: never happened; 1: sometimes; 2: happened regularly), which was categorized into five categories.

### Statistical analyses

For each of the five biological clocks we computed associations with demographic (sex, education), lifestyle (physical activity, smoking, alcohol use), somatic health (BMI, hand grip strength, lung function, physical disability, chronic diseases) and mental health (current depression, depression severity, childhood trauma) determinants using linear models with health determinants as predictors and biological clock as outcome (for each health determinant separately). In all models sex was used as covariate (only not when sex was the outcome). For telomere length, age was used as covariate in the models, for the other biological clocks age was not used as covariate since by design they are independent of age. Correction for multiple testing was done using permutation based FDR^62^. Subject labels were permuted 1000 times and associations were computed using the permuted data (all biological clocks vs all health determinants). For each of the observed *P*-values (*p)* the FDR was computed as the average number of permuted P-values smaller than *p*, divided by the amount of real *P*-values smaller than *p*, resulting in a *P*-value threshold of 2e-2 for a FDR of 5% for all tests. In the 653 overlapping samples with data in each biological clock domain, we scaled (mean 0, standard deviation 1) and summed up the 5 biological clocks in order to create a composite index of biological aging.

## Acknowledgments

The infrastructure for the NESDA study (www.nesda.nl) is funded through the Geestkracht program of the Netherlands Organisation for Health Research and Development (ZonMw, grant number 10-000-1002) and financial contributions by participating universities and mental health care organizations (VU University Medical Center, GGZ inGeest, Leiden University Medical Center, Leiden University, GGZ Rivierduinen, University Medical Center Groningen, University of Groningen, Lentis, GGZ Friesland, GGZ Drenthe, Rob Giel Onderzoekscentrum). Telomere length assaying was supported through a NWO-VICI grant (number 91811602). Methylation sequencing was supported by NIMH grant R01MH099110. Metabolomics data were generated within the framework of the BBMRI Metabolomics Consortium funded by BBMRI-NL, a research infrastructure financed by the Dutch government (NWO, grant nr 184.021.007 and 184033111). Gene expression data were funded by the US National Institute of Mental Health (RC2MH089951).

## Author contributions

L.H., R.J., Y.M., B.P., and J.V. conceived the concept and design of the study. K.A.B. and E.G.J.O. generated the methylation data and commented on the manuscript. R.J. analyzed the data. L.H., R.J., Y.M., B.P., and J.V. wrote the manuscript. All authors participated in the interpretation of the study and reviewed and approved the final manuscript.

## Data availability

Gene expression data used for this study are available at dbGaP, accession number phs000486.v1.p1 (http://www.ncbi.nlm.nih.gov/projects/gap/cgi-bin/study.cgi?study_id=phs000486.v1.p1). Other data sources are not available due to privacy issues (these concern large amount of human subject data).

---

**Table S1.**
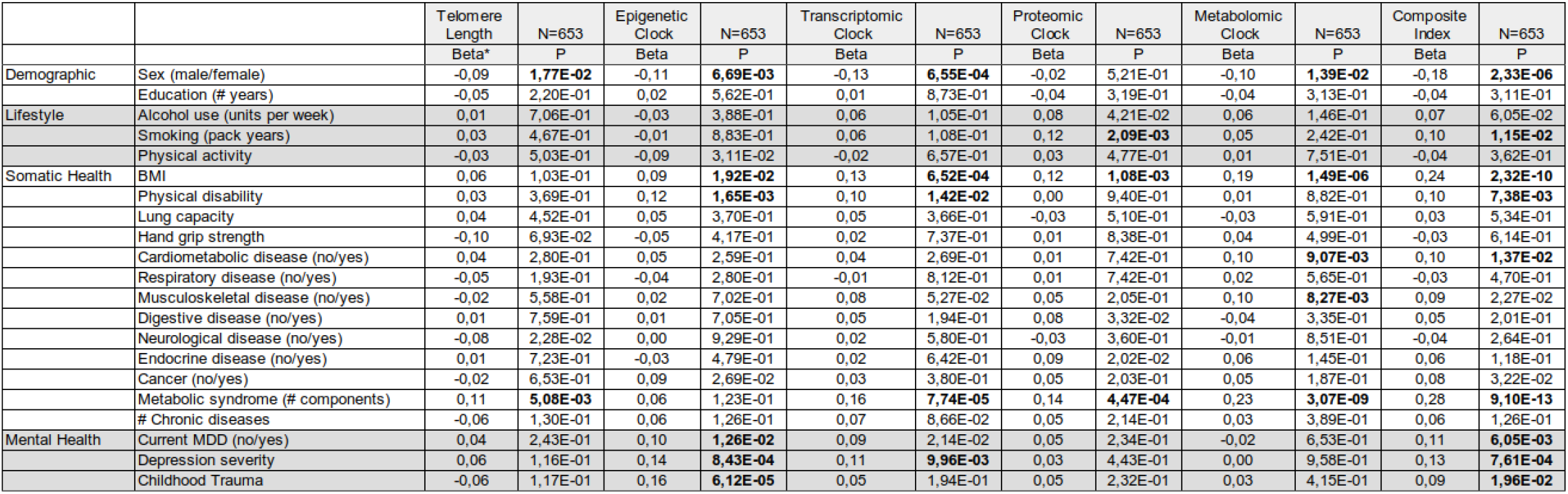
Associations between 5 individual indexes as well as the composite index of biological clocks and multiple health determinants in the 653 overlapping samples. For each biological clock linear models were fit with the health determinant as predictor, while controlling for sex. Analysis was limited to the 653 samples with all 5 data layers available. Beta’s and P-values from these models are presented here.

